# The planktonic microbiome of the Great Barrier Reef

**DOI:** 10.1101/2025.05.13.653689

**Authors:** Steven Robbins, Katherine Dougan, Marko Terzin, Julian Zaugg, Sara C. Bell, Patrick W. Laffy, J. Pamela Engelberts, Nicole S. Webster, Philip Hugenholtz, David G. Bourne, Yun Kit Yeoh

## Abstract

Large genome databases have markedly improved our understanding of marine microorganisms. Although these resources have focused on prokaryotes, genomes from many dominant marine lineages, such as *Pelagibacter* and *Prochlorococcus*, are conspicuously underrepresented. Here, we present the Great Barrier Reef Microbial Genomes Database (GBR-MGD) comprising 5,283 prokaryotic genomes obtained from GBR seawater samples using Nanopore sequencing, including a collection of high quality genomes of underrepresented groups. We show that standard short read assemblies miss these populations due to a combination of strain heterogeneity and low GC% sequencing bias. The GBR-MGD also comprises 20 chromosome-level picoeukaryote and 808,585 viral genomes, including a newly described clade of marine Crassvirales. We demonstrate the use of the GBR-MGD to identify indicator taxa that can reliably predict the effects of reef management practices, such as the establishment of marine protected zones.

## Introduction

Microorganisms dominate life in the oceans^1^, forming the foundation of marine food webs^2^ and driving global biogeochemical cycles^3^ through the metabolic interactions of bacterial, archaeal, viral and microbial eukaryotic communities. Understanding how these communities function, interact, and change over time is critical for predicting ecological resilience, especially as oceans come under increasing pressure from rapid climate change^4^. The overwhelming majority of microbial species found in marine ecosystems have yet to be cultured, though the establishment of large-scale databases of prokaryote genomes via metagenomic sequencing have markedly improved our understanding of the identity, function, and distribution of marine microorganisms^5–7^. Amongst the prokaryotic inventories, however, many of the dominant species driving critical planetary biogeochemical cycles are conspicuously underrepresented, including *Pelagibacter*, *Prochlorococcus*, SAR86, and others^8,9^. Further, these surveys have largely ignored viruses and eukaryotes, with a few notable exceptions^10–12^. Australia’s oceans are also mostly absent in global marine genome surveys^13,14^, as are coral reefs, which are hotspots for biodiversity^4^. To address these gaps, we provide the first detailed assessment of the pelagic microbiome of the world’s largest coral reef, Australia’s Great Barrier Reef (GBR), using hybrid long-read DNA sequencing to construct a holistic database of bacteria, archaea, eukaryotes, viruses, and plasmids, collectively termed the Great Barrier Reef Microbial Genomes Database (GBR-MGD).

We show that there are several prokaryote taxa that cannot be recovered with Illumina short reads alone due to a combination of high strain heterogeneity and a bias against low GC taxa, which can be rectified by inclusion of Nanopore long read sequencing. This technology also enabled us to assemble near-complete Crassvirales genomes that form a distinct group within the order, as well as chromosome-level picoeukaryotic MAGs (eMAGs) from multiple *Bathycoccus* structural variants and the as-yet undescribed *Ostreococcus* Clade B, often the most common picoeukaryote found on coral reefs. We were able to leverage the GBR-MGD prokaryote MAGs (pMAGs) and machine learning techniques to identify reproducible shifts in community structure that reflect the Great Barrier Reef Marine Park Authority’s management plan designating reefs as open or closed to commercial fishing. To our knowledge, this is the first demonstration of an ecosystem-wide effect of fisheries management on the microbial communities of the surrounding seawater. This study marks a substantial advance in our ability to recover complete and near-complete prokaryotic, eukaryotic, and viral genomes from seawater samples, enabling marine ecosystem research and management.

## Results

### Sample collection and benchmarking Nanopore hybrid assemblies

The absence of representative microbial genomic surveys of Australia’s Great Barrier Reef hampers our ability to understand how reef microbial communities are changing, especially in response to anthropogenic stressors such as climate change. To generate Australia’s first comprehensive database of marine microorganisms, seawater samples were collected from 48 sites spanning a north to south transect of the GBR (**Fig. 1**) for metagenomic sequencing (**Tables S1-3**). To enhance MAG recovery, we used a hybrid assembly of long and short reads, which has been shown to substantially improve the quality of MAGs compared to short read assemblies^15^.

**Fig. 1:**
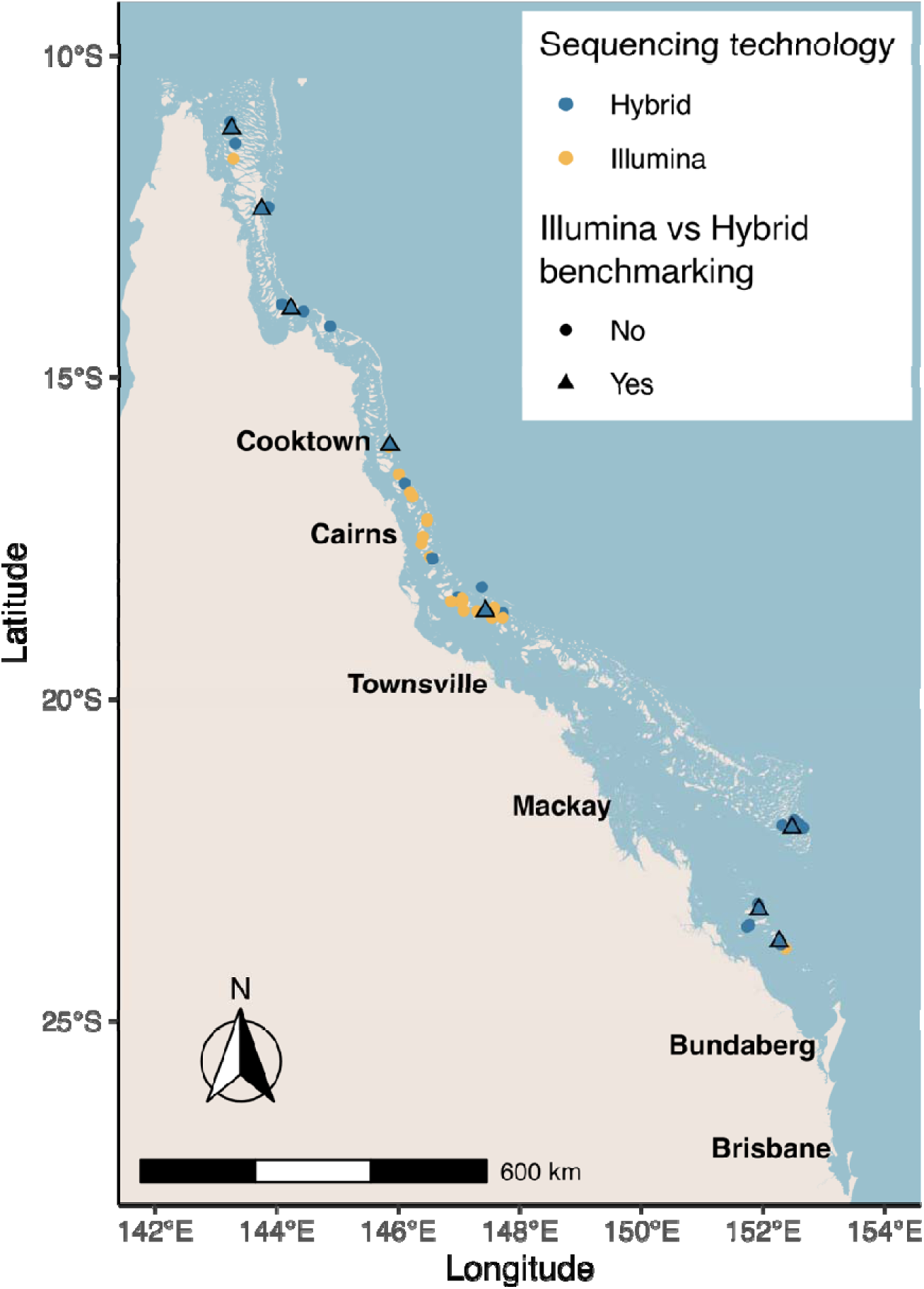
Location of the 48 sites sampled across the Great Barrier Reef. Sampling was conducted alongside physicochemical surveys performed by the Australian Institute of Marine Science’s (AIMS) Long Term Monitoring Program (LTMP).

We hypothesized that common but difficult to recover marine taxa such as *Pelagibacter* could be obtained using Nanopore long read sequencing to span strain-variable regions based on the expectation of high strain heterogeneity^16^. To test this hypothesis, we selected eight seawater samples for Illumina-only and hybrid (Illumina + Nanopore) assembly, and compared the recovery of medium to high-quality (MQ-HQ) pMAGs with a focus on lineages previously found to be difficult to recover in pMAGs. The total number of base pairs in both Illumina-only and hybrid assemblies was standardized to 30 Gbp depth to remove sequencing depth as a factor. Contiguity of the hybrid-assembled pMAGs was 29-fold higher than that of the Illumina-only pMAGs (N50 of 730 vs 25 kbp), exceeding the genome length of some marine taxa (*e.g.* Nanoarchaeota^17^; **Fig 2A, Tables S4-5, Fig. S1**). Commensurately, the average number of MQ-HQ pMAGs per sample increased ∼2-fold in hybrid assemblies (64 to 123) with the increase even more apparent for HQ pMAGs (13 to 40; **Fig. 2A, Fig. S1**).

**Fig. 2:**
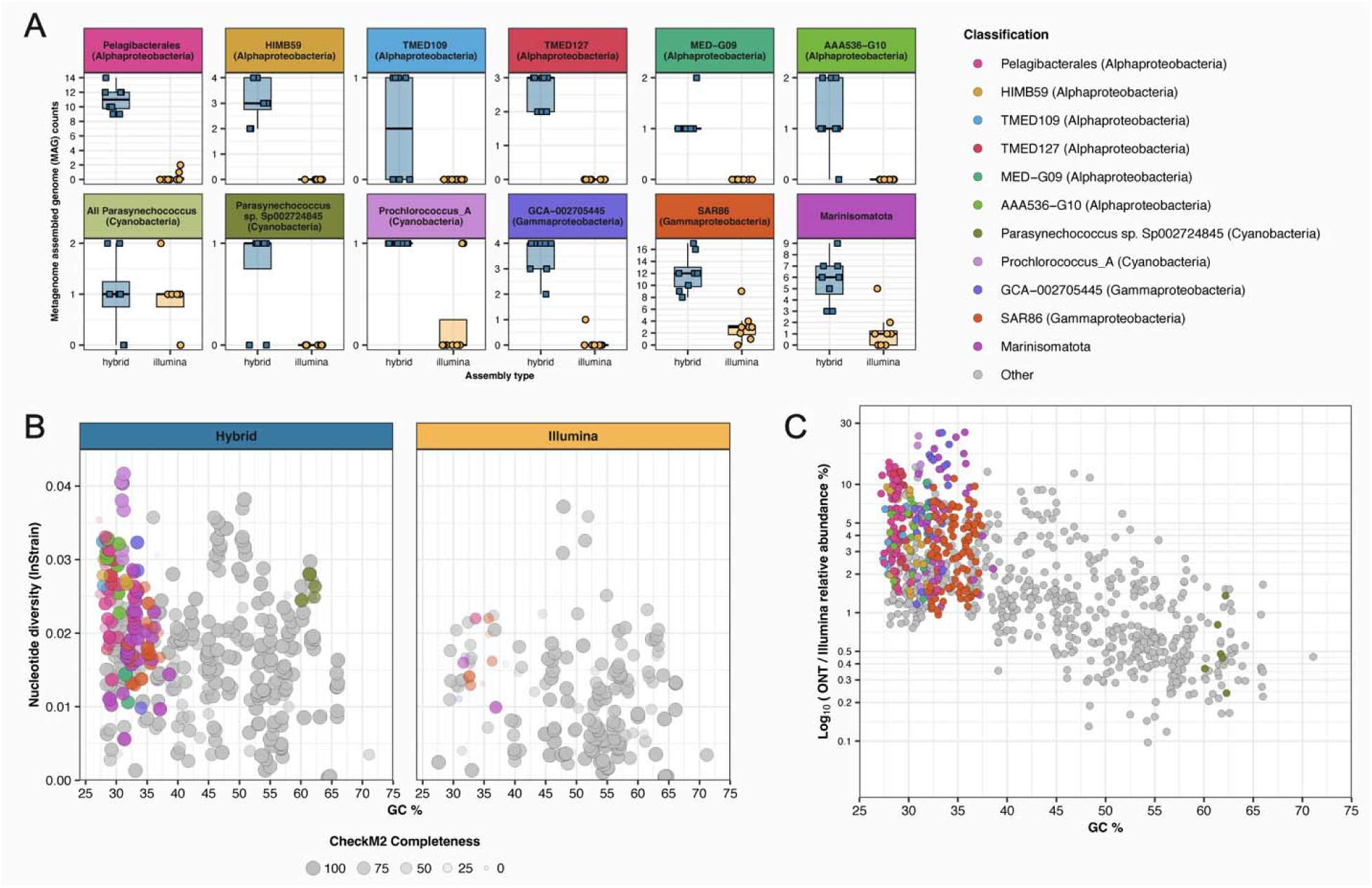
Plots illustrating recovery and sequencing bias between Illumina-only and Nanopore hybrid pMAGs: A) Box and whisker plots showing number of Illumina and Hybrid pMAGs recovered from taxa showing substantial underrepresentation in Illumina-only benchmark pMAGs. B) plots of Nucleotide Diversity and average GC% for Hybrid (left panel; N=382) and Illumina-only (right panel; N=179) benchmark pMAGs after species-level dereplication (95% ANI), highlighting taxa shown to be challenging to recover using short reads in plot A. Circle opacity and size are scaled according to CheckM2 completeness. C) plot showing Nanopore overrepresentation as a function of average MAG GC% across all benchmark Hybrid (N= 987) and Illumina-only (N=516) pMAGs, where overrepresentation is defined as the ratio of pMAG relative abundance in Nanopore vs Illumina reads. All MAGs have CheckM2 quality scores >50 and contamination <10%. Circle color in panels B and C correspond to those used in panel A.

The increase in the number of overall hybrid pMAGs included lineages previously identified as difficult to recover with short read-only sequencing, including Pelagibacterales, *Parasynechococcus*, *Prochlorococcus* and SAR86, along with other lineages not previously recognized as difficult to recover. For example, pMAGs from the Alphaproteobacteria classes HIMB59, TMED109, TMED127, MEDG09, the family AAA536-G10 within the order Puniceispirillales (formerly SAR116), and the Gammaproteobacteria order GCA-002705445, were not recovered from Illumina-only assemblies but were consistently recovered in hybrid assemblies (**Fig. 2A**). Strikingly, genomes belonging to the order Pelagibacterales and phylum Marinisomatota (formerly Marinimicrobia / SAR406) were recovered infrequently in Illumina-only assemblies, whereas 5 to 17 pMAGs were recovered per sample from the hybrid assemblies (**Fig. 2A**), resulting in substantial missed diversity using short reads only assemblies—25 *vs* 2 different species across the eight samples for Pelagibacterales and 13 *vs* 7 species in Marinisomatota (**Tables S4-5**). Within the Cyanobacteriota, *Prochlorococcus* pMAGs were generally not recovered from Illumina-only assemblies but were recovered from all hybrid assemblies. Interestingly, while *Parasynechococcus* (formerly *Synechococcus_E*) pMAGs could be recovered from both short read and hybrid assemblies, only the hybrid assemblies recovered representatives of *Parasynechococcus* species sp002724845 (**Fig. 2A**), the most dominant prokaryote lineage in GBR seawater, averaging 8.9 ± 5.9% relative abundance (max 35.9%) across the complete GBR-MGD dataset. By comparison, the next most abundant *Parasynechococcus* species averaged only 0.2% (**Table S6**).

We next investigated whether strain heterogeneity was the driving factor behind the improved recovery of pMAGs using Nanopore hybrid assemblies. One way to assess strain heterogeneity is to quantify single nucleotide polymorphism (SNP) frequencies, and indeed, a high proportion of hybrid pMAGs from difficult-to-recover taxa represent populations with high nucleotide diversity (>0.02, **Fig. 2B**), a measure of SNP frequency within the metagenomic reads^18^. By contrast, Illumina-only pMAGs rarely have nucleotide diversity values >0.023, indicating a threshold above which short-read pMAG recovery is severely impacted (**Fig. 2B**). However, several difficult-to-recover taxa have sufficiently low strain heterogeneity (0.005 to 0.02; **Fig. 2B**) to be recovered by Illumina-only assembly, suggesting the possibility of a secondary reason for improved pMAG recovery in hybrid assemblies. By mapping both the Illumina and Nanopore reads used in hybrid assemblies back to their respective pMAGs, we found that difficult-to-recover taxa were consistently under-represented in Illumina datasets (up to 18x less than in Nanopore datasets; **Fig. 2C**). Plotting GC-content and fold-Nanopore overrepresentation for all pMAGs, defined as the relative abundance in Nanopore reads divided by the relative abundance in Illumina reads, we find a clear and substantial bias in the Illumina reads against pMAGs with low GC content (**Fig. 2C**). For example, Marinisomatota pMAGs are up to 18x less abundant in Illumina reads than Nanopore (avg. 6x), *Prochlorococcus* are up to 16x less abundant (avg. 7x), Gammaproteobacteria GCA-002705445 are up to 15x (avg. 6x) less abundant, and *Pelagibacter* are up to 7x (avg. 3x) less abundant. In addition to high nucleotide diversity, this would account for the inability of Illumina-only assemblies to recover pMAGs from difficult taxa, as all have GC contents below 40%. This is consistent with a known GC bias in polymerase-based sequencing platforms, including Illumina^19^. When considering both GC bias and strain diversity as factors, short read MAG recovery appears to be difficult above 0.005 strain diversity and less than 40% GC, as few pMAGs were recovered within this range and those that were showed lower completeness values (**Fig. 2B**). The problem reaches a critical threshold above 0.023 strain diversity and <40% GC, where no short read pMAGs could be recovered, despite many difficult taxa showing values in this range and being readily recovered in hybrid pMAGs (**Fig. 2B**). In summary, we show that assemblies incorporating Nanopore long reads are key to recovery of dominant and ubiquitous marine lineages missed by short read-only assemblies, not only because long reads can overcome elevated strain heterogeneity but because of GC bias inherent to sequencing-by-synthesis platforms.

### Recovery of prokaryote MAGs from GBR-MGD seawater

Having established that Nanopore sequencing is essential for representation of cosmopolitan marine lineages, we extended our efforts to additional samples to produce the Great Barrier Reef Microbial Genomes Database, or GBR-MGD. Hybrid assemblies from 27 of the 48 GBR seawater samples (**Fig. 1**) yielded 4,713 bacterial and archaeal pMAGs, including 1,505 high-quality (>90% completeness with <5% contamination) and 345 circular pMAGs assumed to be complete. We supplemented these data with Illumina-only assemblies from the remaining 21 sites resulting in an additional 570 pMAGs (85 HQ), bringing the total number of bacterial and archaeal pMAGs to 5,283 in the final GBR-MGD database. Of the taxa identified to be recalcitrant to pMAG recovery, we obtained 319 Pelagibacterales pMAGs (22 HQ, 7 circular), 445 SAR86 (128 HQ, 35 circular), 222 Marinisomatota (70 HQ, 14 circular), 21 *Parasynechococcus* sp002724845 (1 HQ, 0 circular), 26 *Prochlorococcus* (8 HQ, 0 circular), 135 Gammaproteobacteria order GCA-002705445 (67 HQ, 34 circular), and 45 Alphaproteobacteria family AAA536-G10 (4 HQ, 0 circular). The recovery of circular pMAGs also allowed us to identify lineages for which CheckM version 1 (CheckM1) systematically underestimates completeness, as circular pMAGs should technically have a completeness value of 100%. Indeed, all circular genomes from the following taxa had completeness estimates below 85%: Gammaproteobacteria orders SAR86 and GCA-002705445, Bacteroidota genus MED-G13, Chloroflexota order UBA1151, Desulfobacterota class UBA1144, and the archaeal order Poseidonales (**Table S7: GBR-MGD MAG stats**). Completeness estimates were substantially higher for circular genomes using CheckM2, averaging 91% *vs* 79% for CheckM1, though groups GCA-002705445 and SAR86 are still chronically underestimated using CheckM2 (avg ∼75% completeness) despite being circular. This underscores the need to use CheckM2 for completeness estimates of marine pMAGs to avoid discarding systematically underestimated genomes, which has contributed to underrepresentation of certain taxa^8,9^.

Species-level dereplication (≥95% ANI) of the final set of GBR-MGD pMAGs resulted in a dereplicated set of 876 unique species. To determine the novelty of the GBR-MGD data, we compared it to the ∼35,000 pMAGs present in the Ocean Microbiomics Database^13^ (OMD), revealing that 65% of the GBR-MGD species were not previously represented (<95% ANI to OMD sequences). These may therefore represent either newly recognized marine species potentially specific to the GBR or taxa that are present elsewhere but are not represented in public databases due to difficulties with assembly. Notably, of the 327 pMAGs (35%) that are part of a previously identified species cluster in the OMD, GBR-MGD pMAGs were the best representative of the cluster two thirds of the time based on CoverM Cluster scoring criteria, which takes into account genome completeness, contamination, and number of contigs in each pMAG. Thus, the GBR-MGD comprises a wealth of new species diversity captured in high quality pMAGs. Mapping of the GBR-MGD Illumina reads from all samples to the dereplicated pMAGs (**Table S6**) showed that an average 47 ± 11% (max 66%) of the community was represented by prokaryotic taxa. This result exceeds previous read-mapping statistics of marine metagenomes against large prokaryote genome databases, including TARA oceans (2,631 MAGs; 7% average mapping)^5^, GEMS^13,20^ (8,578 MAGs; ∼20% average mapping), SAG-only GORG Tropics Database^13,21^ (20,288 SAGs; ∼20-40% average mapping), and is nearly on par with the OMD, a gold standard aggregated set of >35,000 MAGs and SAGs^13^ from all previous marine studies (∼58% average mapping for the 0.2-3 µm size fraction).

### Potential of prokaryote MAGs as indicators of reef protection status

Marine protected areas are refuge zones created to protect sensitive reef-dwelling animals from human disturbance. When strictly enforced, such areas are known to be effective tools for reef conservation that can buffer the negative effects of commercial fishing and marine heatwaves on sensitive species^22^. Consequently, the GBR Marine Park Authority has categorized reefs into zones open to fishing (Fished) and No-Take Marine Reserves (NTMRs) that are closed to fishing to protect ecosystems and maintain stable populations of commercially harvested fish (**Fig. S2 and Table S8**). Marine microbial communities have been shown to shift rapidly and predictably in response to environmental change such as nutrient loads and temperature, which could be leveraged to inform management decisions^23^. To test whether seawater microbial communities can predict reef protection status, we analyzed the abundance of pMAGs from co-located Fished and NTMR reefs (**Fig. S3**) using a supervised machine learning classifier (MINT sPLS-DA^24^) and were able to discriminate (**Fig. 3A**) and predict protection status with 71% accuracy, suggesting an ecosystem-wide effect that extends into the surrounding seawater to influence microbial communities. Interestingly, the majority of pMAGs indicative of NTMR status belong to Pelagibacterales, SAR86, TMED109, HIMB59, and Marinisomatota, all identified in the present study as lineages that are recalcitrant to recovery using short read sequencing due to high strain heterogeneity and low GC content (**Fig. 2**). Many of these taxa have undergone genome streamlining (reduced cell and genome size and lower GC content; **Figs. 3B, C, and E**)^25–27^ as an adaptation to oligotrophic habitats to reduce their requirement for nutrients. By contrast, the majority of taxa indicative of Fished reefs have GC contents >45% with no evidence of genome streamlining (**Figs. 3B, C, and E**), including members of the genera UBA11663, UBA8752, and species UBA10364 sp003445735 in the class Bacteroidia, the family Halieaceae in the Gammaproteobacteria, and the species HIMB11 sp000472185 in the Alphaproteobacteria (**Figs. 3A, D, and E**). In fact, at ∼60%, 63%, and 45% average GC respectively, the UBA11663, UBA8752, and UBA10364 pMAGs have GC contents 9 to 27% higher than the average member of the Bacteroidia (36%) from the GBR-MGD (**Table S7**).

**Fig. 3:**
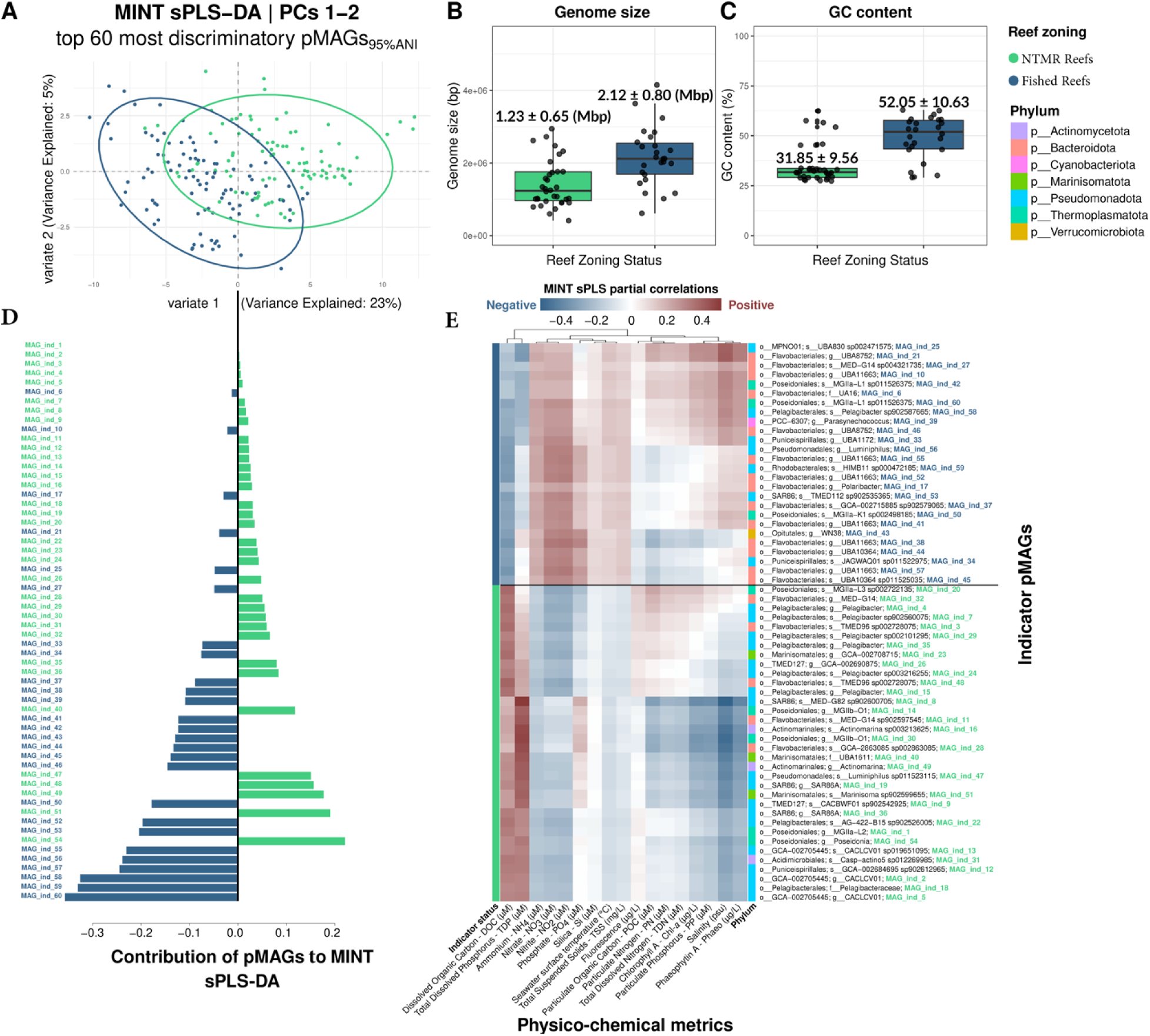
Microbial indicators of reef protection status. A) MINT sPLS-DA plot showing clustering of samples by Fished vs NTMR status, showing that the top 60 most influential indicator pMAGs can discriminate Fished vs NTMR reefs. B) Average genome size of the indicator pMAGs (95% ANI dereplicated) from Fished and NTMR reefs. C) Average GC content of pMAGs from Fished and NTMR reefs. D) Bar chart showing loading weights of all 60 indicator pMAGs ranked from most influential (bottom) to least influential (top) in discriminating Fished vs NTMR reefs, selected from MINT sPLS-DA dimension 1. E) Heatmap of MINT sPLS partial correlation coefficient values for each indicator pMAG showing positive (red) or negative (blue) correlation with seawater physicochemical variables. Indicator MAG taxonomy is listed to the right of panel E and includes a MAG ID in the form MAG_ind_# that maps to the same entry in panel D.

The association of NTMR reefs with streamlined, low-GC taxa that require less nitrogen for DNA synthesis^26–29^ than their higher-GC counterparts on Fished reefs suggests that protection status may influence reef nutrient levels. Leveraging nutrient measurement data taken alongside our microbial samples as part of the Australian Institute of Marine Science’s (AIMS) Long Term Monitoring Program (LTMP), we find that dissolved nitrogen (ammonia, nitrate, and nitrite) is indeed lower in reefs closed to fishing than open reefs (**Fig. S4**), suggesting that differences in nutrient levels between Fished and NTMR reefs select for taxa with different ecological and evolutionary strategies. An increase in nutrient loads with decreased fish density has been previously demonstrated and may account for higher nutrient levels in Fished reefs^30–32^. The inability of short-read sequencing to deliver pMAGs from streamlined, low-GC taxa, which are ubiquitous in marine ecosystems, has not previously been appreciated and highlights that Nanopore sequencing was necessary to identify these patterns.

### Recovery of GBR-MGD DNA viruses

Viruses are the most abundant biological entities in marine environments, strongly impacting marine ecology through interactions with their prokaryote and eukaryote hosts. For example, it is estimated that viruses kill 20% of marine microbial biomass per day, representing a substantial carbon turnover^33^. Viral detection tools perform poorly on highly fragmented assemblies with shorter contigs, and consequently produce variable results^34,35^. Therefore, we first tested six widely used metagenomic viral prediction tools to identify viruses in the GBR-MGD; GeNomad^34^, Virsorter2^35^, ViralVerify^36^, VIBRANT^37^, PPR-Meta^38^, and DeepVirFinder^39^, using CheckV to assess the number of complete viruses predicted by each tool. We further plotted the ratio of host (*i.e.* prokaryote) to viral marker genes identified by CheckV within each putative viral contig to identify spurious assignments. DeepVirFinder showed substantially more false positives than any other tool, with the proportion of bacterial marker genes among putative viral contigs increasing with contig length, indicating clear bacterial origin (**Fig. S5 and S6**). Surprisingly, we also found that CheckV showed a similar fault, where long contigs were increasingly likely to be designated as complete viral genomes, despite having a high ratio of bacterial to viral markers (**Fig. S5 and S6**). We therefore excluded DeepVirFinder from our pipeline as unsuitable for use with long-read metagenomes and the remaining five tools were used in concert to produce a consolidated dataset of 808,585 single contig viral MAGs (vMAGs) from across the 48 GBR-MGD reefs, representing 362,802 species-level (95% ANI) vOTUs (**Table S9**). Read mapping against the vOTU representatives accounted for between 4 to 27% of each metagenome (averaging 13 ± 4%), highlighting that viruses comprise a significant proportion of GBR seawater metagenomes^33,40^.

The majority of vOTUs were classified as Caudoviricetes (67% of sequences), of which only 6% were resolved below this class. This included the families Kyanoviridae (2.8%) encompassing the T4-like cyanophages that commonly infect *Prochlorococcus* and *Synechococcus*^41^; Autographiviridae (1.4%), common phages of *Prochlorococcus*, *Synechococcus,* and *Pelagibacter*^42–45^; and the order Crassvirales (1%; see below). A further 6% of the vOTUs were classified as giant phages belonging to the Nucleocytoviricota, which likely reflects our ability to recover low GC genomes characteristic of this phylum. The remaining vOTUs (27%) were unclassified, which may indicate *bona fide* marine viral novelty or difficulties in discriminating viruses from other genetic elements. Therefore, we provide the CheckV and GeNomad marker gene frequency tables to allow users to filter vMAGs based on their own criteria (**Tables S10-12**).

### Recovery and analysis of marine Crassvirales

Crassphages were first identified in cross assemblies of human gut metagenomes^46^ and are currently recognized as the most abundant phage in the human gut^47,48^. Although members of the Crassvirales have since been identified in a range of additional environments such as primate and termite gut samples, terrestrial systems, groundwater, and marine habitats^49–52^, Crassvirales from these habitats remain poorly characterized. Within the GBR-MGD, 312 vMAGs >20 kb in length were classified by GeNomad as Crassvirales, of which 167 contained a terminase large subunit marker gene (*terL*) containing at least 400 amino acid residues. However, phylogenetic placement within a TerL tree containing representative genomes from the class Caudoviricetes, indicated that the majority of these sequences (155, 93%) are actually members of sister clades to the Crassvirales, the families Pachyviridae and Pervagoviridae, which mostly comprise free-living marine flavobacterial phages^53^ (**Fig. 4A**). To confirm the placement of the GBR-MGD Crassvirales within the order, we constructed whole genome phylogenies including all putative Crassvirales GBR-MGD vMAGs >20kb in length and select reference genomes from the Caudovicites, including the Pachyviridae and Pervagoviridae (**Fig. 4B**). This tree also contained recently discovered Crassvirales sequences from Antarctic seawater^54^, providing an opportunity to assess the phylogenetic distribution of the Crassvirales across global oceans and between tropical coral reefs and Antarctic waters (**Fig. 4B**). Piedade et al. (2024) previously identified four distinct clades of Antarctic Crassvirales, given placeholder names C1 through C4^54^. Notably, GBR-MGD Crassvirales are part of a larger monophyletic clade of exclusively marine sequences that includes the C1 and C2 clades (**Fig. 4A**). We confirm that all members of this marine lineage contain the full complement of core Crassvirales genes^54^ (**Fig. 4C**). Interestingly, most GBR-MGD Crassvirales form a separate clade within the marine lineage characterized by a higher GC content (average 45% *vs* 34%; **Fig. 4B**). However, several cluster with C2 genomes indicating that the C2 clade is not limited to the Antarctic but is also present in Australian tropical reefs, revealing a wider geographical distribution and temperature range (**Fig. 4B**). Similar to animal-associated Crassvirales, the marine lineage, both Antarctic and GBR, are predicted to infect Bacteroidota hosts^31^, suggesting that this association is universal to this viral order. Further work is necessary to delineate the global distribution and ecological characteristics of marine Crassvirales.

**Fig. 4:**
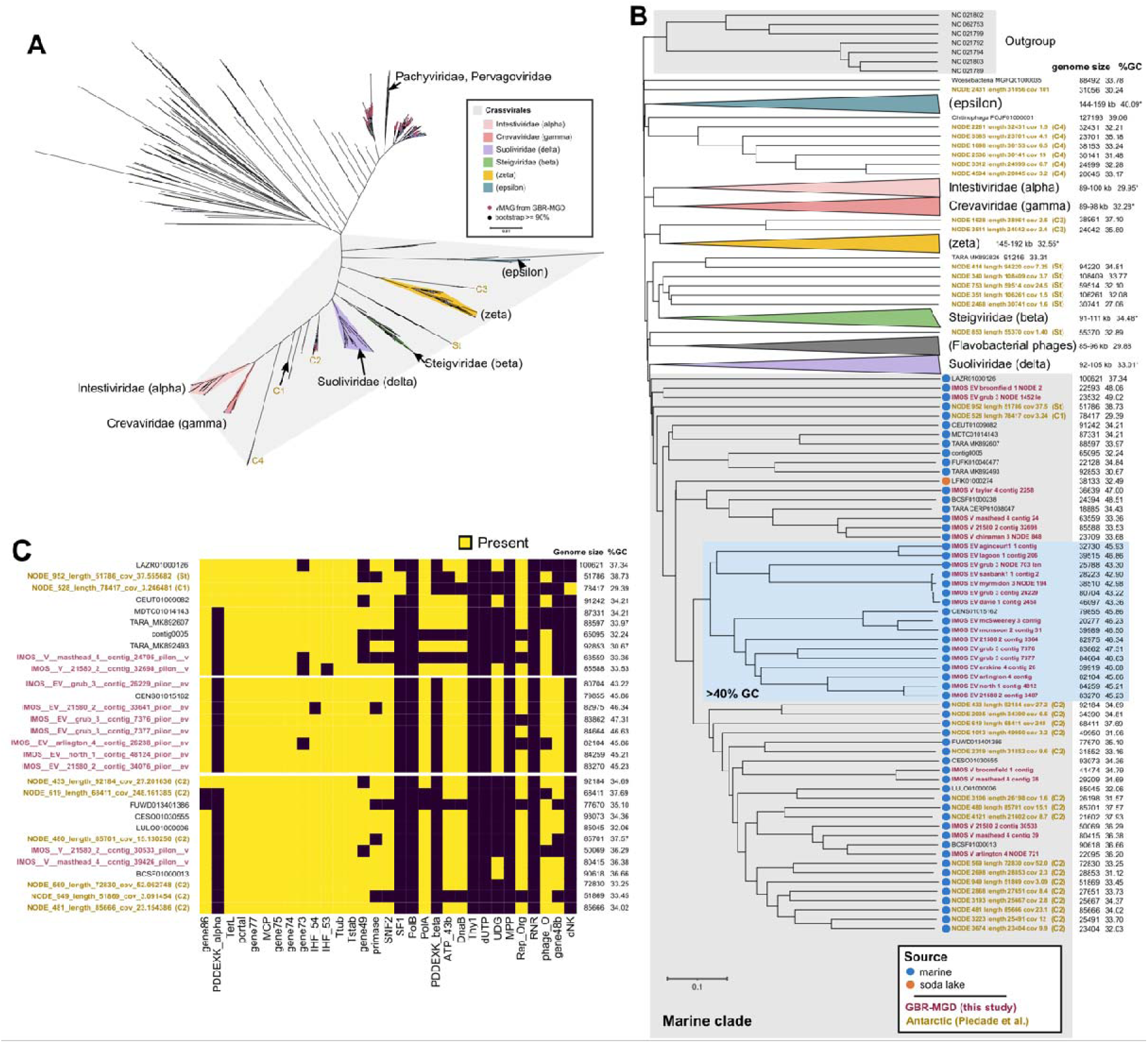
Expansion of the marine Crassvirales. A) TerL tree of Crassvirales and other Caudoviricetes showing placement of MGD Crassvirales vMAGs represented by red circles in relation to existing Crassvirales. Families are labeled with ICTV names along with the previously proposed alpha, beta, gamma, delta, epsilon and zeta group names in parentheses. C1 - C4 and St groups are as proposed in Piedade et al. 2024. B) Whole genome phylogeny (ViPTree) of GBR-MGD Crassvirales vMAGs >20 kb (N = 27). C) Pattern of Crassvirales gene conservation in vMAGs and genomes within the marine clade. Only genomes >50 kb are shown. Genes included here were previously identified in Yutin et al. (2021). gene86: putative structural protein gene 86, PDDEXK: PD-(D/E)XK family nuclease, TerL: terminase large subunit, portal: portal protein, gene77: putative structural protein gene 77, MCP: major capsid protein, gene75: putative structural protein gene 75, gene74: putative structural protein gene 74, gene73: putative structural protein gene 73, IHF_54: IHF subunit gene 54, IHF_53: IHF subunit gene 53, Ttub: Tail tubular protein (P22 gp4-like), Tstab: tail stabilization protein (P22 gp10-like), gene49: uncharacterized protein gene 49, primase: DnaG family primase, SNF2: SNF2 helicase, SF1: SF1 helicase, PolB: DNA polymerase family B, PolA: DNA polymerase family A, ATP_43b: AAA domain ATPase, DnaB: phage replicative helicase DnaB family, Thy1: thymidylate synthase, dUTP: dUTPase, UDG: Uracil-DNA glycosylase, MPP: metallophosphatase, Rep_Org: replisome organizer protein, RNR: ribonucleotide reductase, phage_O: bacteriophage replication protein O, gene48b: phage endonuclease I, dNK: deoxynucleotide monophosphate kinase. Colour labels are consistent across all panels.

### Recovery of chromosome-level picoeukaryote genomes

Marine phytoplankton are responsible for around half of all primary production on the planet^3^, and although picoeukaryotes typically make up a smaller fraction of the phytoplankton community compared to prokaryotes, they exhibit much higher growth and turnover rates and often account for the majority of primary productivity^55,56^. Despite their ubiquity in marine ecosystems^11,57^, much of our knowledge of these groups relies on analysis of a few cultured isolates, none of which are from coral reefs or Australian oceans^58–60^. Researchers have recently begun to recover genomes from uncultured representatives via metagenomics^11^. However, as picoeukaryotes have more complex genomes than their prokaryotic counterparts, including multiple chromosomes, large genome rearrangements, and more extensive repeat regions than prokaryotes, the few available short read-based eukaryote MAGs (eMAGs) are mostly fragmented and incomplete^11^.

Using Nanopore long reads, we were able to obtain telomere-containing (**Table S13**) chromosome-level eMAGs from the Great Barrier Reef for the picoeukaryote genera *Bathycoccus* and *Ostreococcus* (green algae in the order Mamielalles), taxa ubiquitously distributed in the world’s oceans^55^ (**Figs. 5A and B**). In total, five *Bathycoccus* eMAGs were recovered and found to represent two distinct genome variants, BV1 and BV2, with BV1 being closely related to *Bathycoccus prasinos* RCC1105 (96% ANI) and BV2 representing a more distantly related lineage with only 82% ANI to RCC1105, along with a large ∼350 kb inversion on chromosome 4 that is not present in BV1 (**Fig. 5C; Fig. S7-8**). Mapping of the metagenomic reads to representatives of each variant (BV1: reef 11049, BV2: Grub Reef) revealed that they had similar relative abundance across most samples (average 0.07 ± 0.11% for BV1 and 0.08 ± 0.08% for BV2), except for samples collected in winter in which BV2 was ∼7 fold more abundant (average 0.47%, max 2.4%; **Table S13**). Previous marker gene-based studies suggest the existence of multiple *Bathycoccus* ecotypes that may represent different species^57,61^, though the genomic underpinnings of ecotype ecology are unclear and may be clarified through the recovery of additional chromosome-level genomes.

**Fig. 5:**
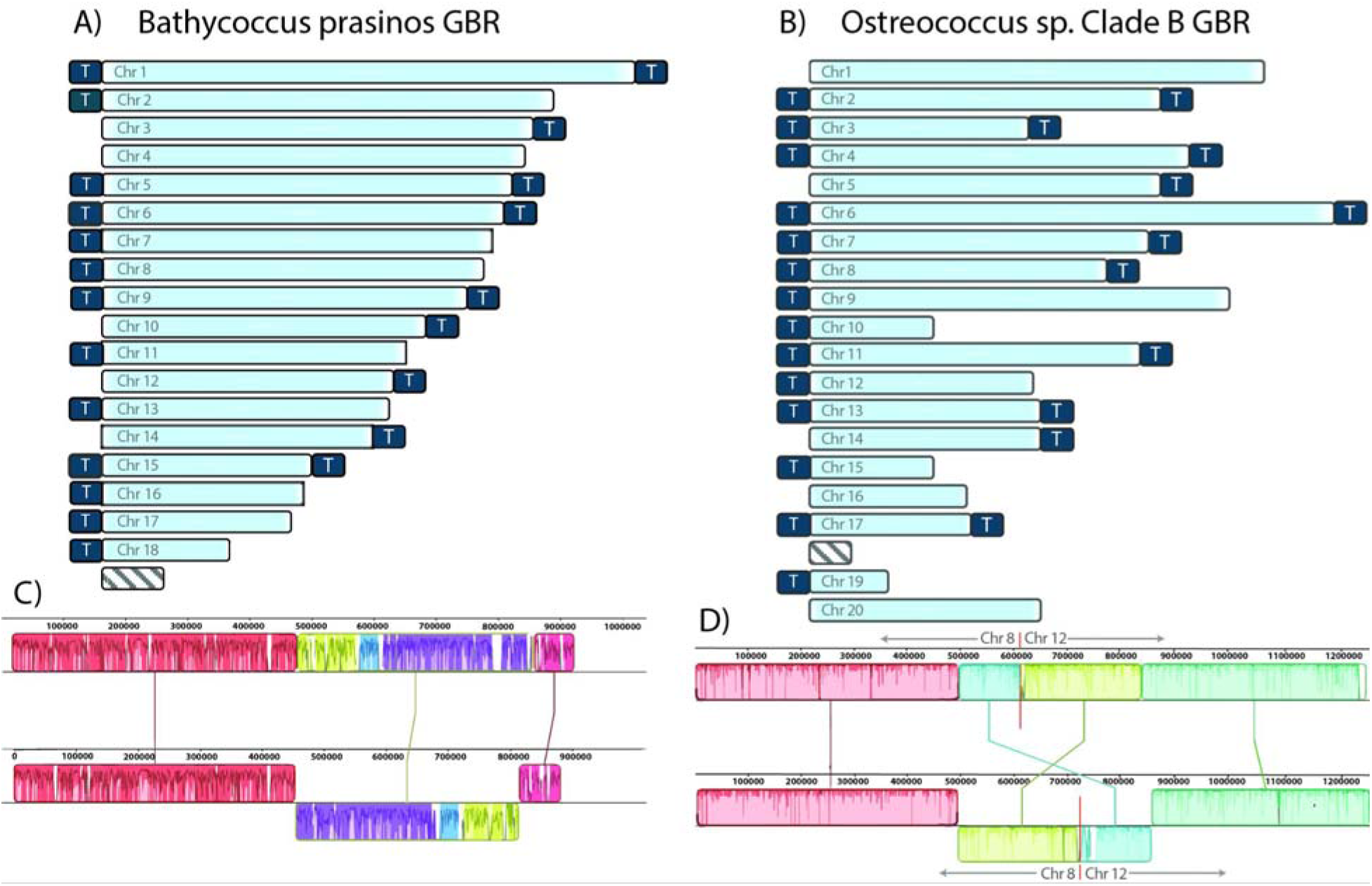
Chromosome-level eMAGs from marine picoeukaryotes. A) Composite illustration of the presence of telomeres in GBR-MGD *Bathycoccus* eMAGs. Presence is noted in dark blue if telomeric repeats were found at the ends of at least one of the five eMAGs, B) Composite illustration of the presence of telomeres in GBR-MGD *Ostreococcus* sp Clade B eMAGs. Presence is noted in dark blue if telomeric repeats were found at the ends of at least one of the 15 eMAGs. C) sequence alignment of chromosome 4 for *Bathycoccus prasinos* RCC1105 (top) and GBR-MGD *Bathycoccus* variant 2 (bottom) from Grub Reef showing a large ∼350kb inversion in variant 2 compared to the reference, D) sequence alignment of chromosomes 8 and 12 from *Ostreococcus* sp. Clade B RCC809 (top) and GBR-MGD *Ostreococcus* sp Clade B (bottom) from Masthead Reef showing a ∼300 kb translocation between the ends of the two chromosomes.

In contrast to *Bathycoccus*, all 15 GBR *Ostreococcus* eMAGs represent a single species closely related to *Ostreococcus* RCC809 (>99% ANI to Ostreococcus RCC809), a member of the Clade B lineage, for which little genomic information is available. Despite high similarity to RCC809, all GBR *Ostreococcus* clade B eMAGs have a ∼350 kb translocation between chromosomes 8 and 12 (**Fig. 5D, Fig. S9**), indicating that high sequence similarity can hide major structural variation detectable via chromosome-level assemblies. After *Parasynechococcus* sp002724845, *Ostreococcus* clade B represents the second most abundant microbial species in the GBR-MGD (average 3.5%, max 30%; **Table S14**). The ability to readily recover eMAGs with long reads, especially to chromosome-level quality, is a substantial advance that we hope will encourage others to explore uncultured eukaryotic dark matter.

## Discussion

Metagenomics has greatly enabled the study of microbial ecosystems by bypassing the cultivation bottleneck. However, marine ecosystems are unusual in that the genomes of many dominant microbial populations have not been recoverable through standard short read metagenomics. Prominent examples include the bacterial genera *Pelagibacter, Prochlorococcus,* and *Synechococcus*, which are all underrepresented in international consortium-wide datasets^8,9^. We show that this phenomenon is due to the inability of short read sequencing and assembly to handle the combination of high strain heterogeneity and low GC content of these taxa. By contrast, the Nanopore platform addresses both issues by being able to sequence low GC templates and to traverse strain variable regions. This enables a representative high quality genome inventory of marine microbial taxa to be generated using readily available technology.

The GBR-MGD will be a highly valuable resource for marine researchers not only because of its inclusion of dominant prokaryotic taxa, but also because it represents a uniquely holistic database linking prokaryote, viral, and eukaryote taxa with long term measurements of nutrient and physicochemical metadata from the AIMS-LTMP. We demonstrate the utility of this connection between the GBR-MGD and LTMP metadata to explain the underlying physicochemical drivers of microbial bioindicators of reef protection status. We were also able to expand the known diversity of marine Crassvirales and note that many of the GBR-MGD sequences form a high GC cluster, which is atypical for this viral order. Finally, we demonstrate that chromosome-level genome recovery is possible for microbial eukaryotes, which comprise up to 30% of pelagic GBR microbial communities.

Moving forward, we suggest that Nanopore long read sequencing should become the standard technology for metagenomic analysis of marine microbial communities, as many key populations are only accessible via this platform. As GC bias is likely inherent to sequencing by synthesis technologies, and strain heterogeneity is not limited to marine environments^18,62–64^, we predict that the sequencing limitations observed in this study will affect metagenomic analyses of other ecosystems that could be addressed through Nanopore sequencing.

## Methods

### Sample collection

Seawater samples were collected from 48 coral reefs spanning the length of the Great Barrier Reef, aligning with *in situ* coral surveys from the LTMP of the Australian Institute of Marine Science in 2019 and 2020 (AIMS; **Fig. 1**). Four 5 L technical replicates of seawater were collected above the reef (2-10 m depth) at each site (N = 192) in either nalgene or Niskin bottles and placed on ice until return to the vessel, where they were filtered successively through 5 µm 33 mm cellulose acetate and 0.22 µm Sterivex-GP filters (Millipore). The Sterivex filters were then snap frozen and stored at −75 °C until DNA extraction.

In addition to metagenomics data, a total of 17 physicochemical (e.g. water chemistry) variables were measured, including ammonia (NHLJLJ), nitrite (NOLJLJ), nitrate (NOLJLJ), total dissolved nitrogen (TDN), phosphate (POLJ³LJ), total dissolved phosphorus (TDP), dissolved organic carbon (DOC), silicate (Si), total suspended solids (TSS), chlorophyll a (Chl-a), phaeophytin a (Phaeo), particulate organic carbon (POC), particulate nitrogen (PN), particulate phosphorus (PP), temperature, salinity, and Chl-a fluorescence (**Fig. 3**) using methods described in Terzin et al. 2025^65^.

### DNA extraction

DNA extractions were carried out inside each 0.22 µm Sterivex filter by adding 1.8 mL filter-sterilised lysis buffer containing 50 mM Tris-HCL (pH 8.0), 40 mM EDTA (pH 8.0), 256 mg/ml sucrose and 18 µl lysozyme (100 mg/ml). Sterivexes were incubated with mild agitation for 1hr at 37°C, followed by the addition of 20 µl proteinase K (20 mg/ml) and further incubation/agitation for 1 hr at 55°C. The lysate was then ejected through the Sterivex into a 5 ml microtube and an equal volume of phenol:chloroform:isoamyl alcohol (IAA; 25:24:1) was added, mixed by inversion and centrifuged at 16,000 x g for 10 minutes. The aqueous phase was then recovered and an equal volume of chloroform:IAA (24:1) was added and mixed by inversion, then centrifuged for 10 minutes at 16,000 x g. DNA was precipitated by adding 1 ml isopropanol and recovered by centrifugation at 20,000 x g for 25 minutes and the supernatant discarded. The DNA was washed with 500 µl 70% ethanol, centrifuged for 10 minutes, the ethanol discarded and the pellet air dried. Finally, 20 µl PCR water was added, incubated at 4°C overnight to resuspend the DNA pellet and then frozen at −20°C.

### Illumina sequencing

All 191 technical replicates from the 48 sites were subjected to Illumina sequencing at the Centre for Microbiome Research (Brisbane, Australia). Libraries were prepared according to the manufacturer’s protocol using Nextera DNA Flex Library Preparation Kit (Illumina # 20018705). Nextera DNA Flex libraries were pooled at equimolar amounts of 2 nM per library. The library pool was quantified in triplicates using the Qubit™ dsDNA HS Assay Kit (Invitrogen) and sequenced using an S4 2×150bp flow cell on a NovaSeq 6000. Additional “deep” sequencing was carried out for one replicate from 27 of the 48 sites to a target depth of 40 Gbp for Nanopore polishing and hybrid assembly (**Table S1-2**).

### Oxford Nanopore sequencing, basecalling, and QC

The replicate chosen for Illumina deep sequencing was also submitted for further long read sequencing on Oxford Nanopore Technology’s (ONT) Promethion platform at the Australian Centre for Ecogenomics (ACE) sequencing facility. For ONT sequencing, DNA was first size selected using the Circulomics SRE XS kit (PacBio, SKU 102-208-200) as this was shown in a small trial to increase total data output and read N50. Native barcoded libraries were prepared from the size-selected DNA following a genomic DNA by ligation protocol (Oxford Nanopore, LSK-109 with EXP-NBD104). Barcoded libraries were pooled with two samples per Promethion flow cell at equal concentration and the library pool was sequenced on a PromethION 24 (Oxford Nanopore) for a total of 72 hours with a R9.4 flow cell using MinKNOW (v20.06.18) with the default settings. We then performed rebasecalling of the fast5 files using the superaccuracy model with Guppy v5.0.16 to generate higher accuracy raw reads before assembly. Finally, Porechop (https://github.com/rrwick/Porechop) was used for barcode trimming using default settings.

### Hybrid assembly

Deep Illumina and nanopore reads for each of the 27 sites were assembled using the Aviary assembly pipeline (https://github.com/rhysnewell/aviary; v0.3.3), specifying the ont_HQ flag for long reads. In brief, by supplying both short and long reads for hybrid assembly, Aviary first assembled the nanopore reads with metaFlye^66^ (v2.9), then polished the metaFlye assembly using a series of racon^67^ (three rounds; v1.4.3), pilon^68^ (one round; v1.23), and racon (one round). Next, a subset of high coverage, polished metaFlye contigs are identified and Illumina reads from the same replicate are mapped to them. Illumina reads that did not map to these contigs were assembled using metaSPAdes^36^ (v3.15.3) together with the nanopore reads. Finally, the hybrid-assembled contigs are collated with the high coverage polished metaFlye contigs to produce the final assembly. Aviary commands were executed in batch using GNU parallel^69^. For the remaining 21 sites not chosen for Nanopore sequencing, a single replicate was assembled using the Aviary pipeline, which uses metaSpades for Illumina-only assembly.

### Prokaryote binning and analysis

Assemblies were binned using the Aviary pipeline’s “recover” workflow, which first mapped reads to each assembly using minimap2 (v2.18). For differential coverage binning, three sites were chosen for mapping to each assembly based on geographical proximity and all nanopore and Illumina reads for all four replicates from each site were used (**Table S15**). Aviary recover then runs binning programs MetaBAT1^70^, metaBAT2^71^ (v2.15), MaxBin2^72^ (v2.2.7), Rosella (https://github.com/rhysnewell/rosella; v0.3.3), CONCOCT^73^ (v1.1.0) and Vamb^74^ (v4.0.0), and includes a bin refinement step through Rosella to further refine bins based on kmer frequency and coverage. Finally, Das Tool^75^ (v1.1.2) was used to collate a representative set of bins from across the different binning tools. pMAGs were retained if they pass a quality threshold ≥50 (quality = completeness - 3 x contamination) using either CheckM1^76^ or CheckM2^77^, such that pMAGs of lower completeness are allowed lower contamination scores for inclusion. Taxonomic classification of the pMAGs was performed using the Genome Taxonomy Database Toolkit (GTDB-Tk^78^) using release R214. Dereplication of MAGs for mapping, and for assessment of presence within previously published databases, was carried out using CoverM^79^ “cluster” at 95% ANI (v0.6). Relative abundance was calculated by mapping the Illumina reads to the dereplicated set of reference pMAGs using CoverM “genome”, requiring 95% identity and 75% read length alignment to consider a read mapped.

### Benchmarks for Illumina vs Nanopore assembly and binning

To directly benchmark the effect of including long nanopore data on improving MAG recovery, we undertook separate re-assembly and binning of eight replicates spanning sites from north to south of the GBR. To remove the effect of sequencing depth, we compared two treatments: 1) Illumina-only assembly of 30 Gbp of sequence data using metaSPAdes and 2) a separate assembly using a subset of 15 Gbp of Illumina reads combined with 15 Gbp of nanopore reads as described above for hybrid assembly. The resulting 16 assemblies were binned with “Aviary recover” to obtain pMAGs as described above.

### Identifying seawater microbes as stable bioindicators of NTMR and Fished reefs

To identify if seawater pMAGs were indicative of NTMR and Fished reefs, we applied Multivariate INTegration Sparse Partial Least Squares Discriminant Analysis (MINT-sPLS-DA^80,81^) using mixOmics^24^ (v6.26.0) to the dereplicated set (95% ANI) of 876 pMAGs (centered log ratio-transformed abundances). MINT sPLS-DA was run in two-steps: tuning the number of MINT sPLS-DA components with *perf()* - to identify that the optimal number of MINT sPLS-DA dimensions (in this case, 1), followed by the *tune()* function to identify the optimal number of pMAGs to retain in the model. This resulted in 260 indicator pMAGs with a minimum balanced error rate of 29% in predicting NTMR vs Fished reefs. We retained the first MINT sPLS-DA component and visualized the top 60 indicator taxa with their loading weights (**Fig. 3**). MINT sPLS-DA results were visualised in mixOmics as 1) the sample plot with the *plotIndiv()* function and as 2) a barchart using *plotLoadings()* which shows the indicator contribution (i.e. loading weights, averaged across GBR sectors) in discriminating NTMRs and Fished reefs (**Fig. 3**). Differences in average genome size and GC content between indicators of NTMRs and Fished reefs were visualized as boxplots in ggplot2 (v3.5.1), and the variation in genome size and GC content was compared between indicators of NTMRs and fished reefs with pairwise Wilcoxon Rank-sum tests in R, which were integrated within boxplots.

### Correlating physicochemical variables with pMAG indicators of reef protection status

To identify environmental drivers that explain the observed patterns between NTMR reefs and Fished reefs, the 60 indicator pMAGs of reef zoning were correlated with each of the 17 physico-chemical variables using MINT sPLS. sPLS^82,83^ models the relationship between multiple predictors (physicochemical variables) and multiple responses (microbial taxa), while MINT^80^ integrates samples from independent subsets to account for confounding effects, such as season and geography. Median values for each physicochemical variable were first calculated per reef site, as the number of replicates differed for molecular (N = 4) and water chemistry (N = 3) samples. We then correlated CLR-transformed abundances of the 876 dereplicated pMAGs with the 17 physicochemical measurements, but have only extracted correlations for the 60 pMAGs that were indicative of NTMR and Fished reefs as our taxa of interest. Similarity scores (partial correlations) were visualized in ggplot2 (v3.5.1). Final composite plots were made in Inkscape (0.92.5).

### Viral, Plasmid, and Eukaryote contig identification

The following viral prediction tools were used to identify putative viral contigs from the GBR-MGD metagenomic assemblies: DeepVirFinder (not versioned; retrieved from GitHub on 8 March, 2023; p-value < 0.05), GeNomad (v1.5.018; --enable-score-calibration --min-score=0.7 --max-score=0.1) with database v1.2, PPR-Meta (v1.134; -t 0.9), VIBRANT (v1.2.132), ViralVerify (v1.133; -thr 7), and VirSorter2 (v2.2.431; --include-groups “dsDNAphage,NCLDV,RNA,ssDNA,lavidaviridae”, --min-score=0.9) with database v1.2. Putative viral contigs were then input to CheckV (v1.0.1) with database v1.5 for quality trimming and assessment. Viral predictions were examined using CheckV’s curated marker sets by visualizing the ratio of host:viral markers and the proportion of viral markers for each putative virus prediction to assess whether the contigs predicted by each tool were host or virus-derived.

In addition to three viral prediction tools that also predict plasmids (GeNomad, PPRMeta, ViralVerify), two additional plasmid-only predictors were used on the metagenomic assemblies, including PlasForest (v1.2), and PlasX (unversioned; score ≥ 0.9) which uses annotated gene families generated in anvi’o (v7.1) using the COG2014 and Pfam v32 databases. Whokaryote (v1.1.2) and EukRep (v0.6.6) were used for eukaryotic prediction.

### Manual curation of picoeukaryote genomes

Genomes for two picoeukaryotes from the *Mamiellales* family, *Bathycoccus prasinos* and *Ostreococcus* sp. Clade B, were manually curated after initial blast results of partial eMAGs revealed the presence of potential *Bathycoccus* and *Ostreococcus* long contigs in high abundance as determined by read mapping. Therefore, complete genomes for the sequenced *Mamiellales* isolates *Bathycoccus prasinos* RCC1105, *Micromonas commoda* RCC299, *Micromonas pusilla* CCMP1545, *Ostreococcus lucimarinus* CCE9901, *Ostreococcus tauri*, and *Ostreococcus* spp. RCC809 were retrieved and used as references for competitive mapping of large metagenomic contigs (≥50 kb) with minimap2^84^ (v2.28) using its genome/assembly alignment mode, allowing up to 20% sequence divergence (--asm20). The majority of contigs were recruited by either *B. prasinos* RCC1105 or *Ostreoccoccus* spp. RCC809, and were selected for manual curation of two distinct species using the RCC1105 and RCC809 genomes as references. Contigs recruited by *Ostreococcus* spp.RCC809 also showed high recruitment to *Ostreococcus lucimarinus* CCE9901, which was therefore used as a secondary reference as needed. Contigs were aligned to the corresponding reference using progressiveMauve from the Mauve aligner^85^. In the few cases where a chromosome was recovered in a sample fragmented in 2-4 contigs, Mauve Contig Mover (MCM) from the Mauve program was used to reorder and link the chromosomal fragments with a linker of 100 N’s. Telomeres were identified at the ends of chromosomal contigs using Tidk^86^ (v0.2.65) to search for telomeric repeat sequence AACCCT, followed by visual inspection to confirm elevated numbers of telomeric repeats at the ends of contigs using (Tidk Plot).

### Final GBR-MGD database compilation

The GBR-MGD database is composed of prokaryotic MAGs, manually curated Mamiellales genomes, and contigs identified as plasmid, viral, or eukaryotic. A standardized labelling scheme with the structure of IMOS__<TaxaGroup>__SAMPLENAME__CONTIG:<TaxaComponent> was applied to all sequences. TaxaGroup indicates all taxa that were linked to that contig (E=eukaryote, M=plasmid/mobile element, P=prokaryote, V=virus), with the presence of multiple letters indicating contigs linked to more than one group. For example, PV would indicate that a contig was present in a pMAG and was also predicted to be a virus or provirus, MPV would indicate that a contig was found in a pMAG and was also predicted to be both a plasmid and a virus, V alone would indicate that the contig was identified to be a virus but is not found in another group. In contrast, contigs marked by only one group were only identified to be in that group. Because viruses can be predicted within other taxa (e.g. PV contigs), TaxaComponent indicates the taxa represented by a particular sequence as well as the bp position in the case of some viral sequences. For example, a proviral region identified within a prokaryotic bin would be represented by two distinct sequences. The sequence within the prokaryotic bin would be labelled IMOS__PV__SAMPLE__CONTIG:p while its counterpart representing the viral prediction would be labelled IMOS__PV__SAMPLE__CONTIG:v1001-10000.

The prokaryotic bins and manually curated Mamiellales genomes were dereplicated together using CoverM. Contig predictions were dereplicated separately for each taxon group (e.g. V, P, E, PV, etc) using pairwise ANI (95% ANI + 85% alignment fraction) using the anicalc.py and aniclust.py scripts provided by CheckV. Relative abundance of each component of the database was calculated by mapping the Illumina metagenomic reads to each taxon group in the final dereplicated database separately using CoverM Genome, requiring 95% identity and 75% alignment to be considered mapped.

### Marine Crassvirales

vMAGS classified by GeNomad as *Crassvirales* were annotated using Pharokka^87^ (v1.6.1) with additional *Crassvirales* protein profiles provided by Yutin et al. 2021^51^. To verify the GeNomad classifications, we downloaded the set of Caudoviricetes TerL sequences compiled by Piedade et al (2024)^54^ and aligned TerL sequences detected in the vMAGs (> 400 amino acid residues) to this collection using MAFFT^88^ (v7.508). The output sequence alignment was trimmed with trimAl^89^ (v1.4.rev15) and used to infer a bootstrapped phylogenetic tree using IQ-TREE2^90^ (v2.2.0.3). Separately, we constructed a proteomic tree using the vMAGs (>20 kb) based on genome-wide sequence similarities computed using tBLASTx as implemented in ViPTree^91^. Both trees were visualised and edited for clarity in ITOL^92^ v7 and Inkscape. Host prediction was performed on vMAGs using iPHoP^93^ (v1.3.3) with database Aug_2023_pub_rw including pMAGs dereplicated at 99% ANI added to this host database.

## Acknowledgements

The seawater samples in this study were collected across 48 reefs from the sea country of various Indigenous groups who are Traditional Owners (TOs) of that land. We acknowledge the Gudang Yadhaigana TOs, custodians of the McSweeney, Monsoon, 11-049, and 11-162 reefs, which lie within their sea country estate. We pay our respects to the Kuuku Ya’u TOs of the Mantis and Lagoon reefs, and the Lama Lama TOs of the 13-124, Davie, and Corbett reefs in the western half of their territory. We also recognize the Cape Melville, Howicks, and Flinders Island TOs of the eastern half of Corbett and Sand bank #1 reefs, as well as the Eastern Kuku Yalanji TOs of St Crispin and Agincourt #1. We extend our respect to the Yirrgandji TOs of Hastings reef and the Gunggandji as TOs of Arlington, Thetford, and Moore reefs. We acknowledge the Gunggandji-Mandingalbay Yidinji TOs of McCulloch, Hedley, Peart, and Feather reefs, and the Mandubarra TOs of Farquaharson Reef. We honour the Girringun Aboriginal Corporation TUMRA for their connection to Taylor Reef, and the Manbarra TOs of Rib, Kelso, Little Kelso, and John Brewer reefs. We also recognize the Wulgurukaba TOs of Myrmidon, Grub, and Helix reefs, the Bindal TOs of Knife, Fork, Centipede, Chicken, and Lynchs reefs, and the continuing connection of the Manbarra TOs to Roxburgh, and Fore and Aft reefs. Lastly, we acknowledge the PCCC TUMRA for their stewardship of North, Bloomfield, Eskine, Mast Head, Hoskyn, Fairfax, and Boult reefs. We pay our respects to their Elders, past, present, and emerging, and acknowledge their enduring connection to land and sea. Further, our desktop / lab research took place at the Australian Institute of Marine Science (AIMS) headquarters at Cape Ferguson, and we wish to acknowledge the Wulgurukaba and Bindal peoples as the TOs of that land. This research was also undertaken at the JCU Townsville Bebegu Yumba campus, and the authors acknowledge that the Australian Aboriginal and Torres Strait Islander peoples are the original inhabitants and traditional custodians of this continent and have unique cultural and spiritual relationships to the land and waters. We acknowledge the AIMS Water Quality team, especially Renee Gruber, Ulysse Bove, Keeley Glasson, and Daniel Moran for logistics, training, and processing of water chemistry samples. We acknowledge the AIMS-LTMP team and others involved in field collection and preparation of samples including Michael Emslie, Emmanuelle Botté, Johnston Davidson, Véronique Mocellin, and Josephine Nielsen. We thank the crew of the RV Solander and RV Cape Ferguson for their excellent logistical support in the field.

## Declarations

## Ethics approval and consent to participate

Samples were collected under the permit G12/35236–1 issued by the Great Barrier Reef Marine Park Authority.

## Consent for publication

Not applicable.

## Data availability

The 5,283 pMAGs, 15 Ostreococcus Clade B genomes, 5 Bathycoccus genomes, and 27 Crassvirales genome constituting the GBR-MGD are in the process of being uploaded to ENA under the project accession XXXXXXXXX and secondary accession XXXXXXXX under the project name “Great Barrier Reef seawater microbiomes genome database. We affirm that these data will be publicly available before publication.

## Author Contributions

NSW obtained funding for the project. NSW and SJR conceived the sampling design. SCB collected seawater in the field and processed all samples in the laboratory for metagenomic sequencing. SJR, MT, YKY, KD, JPE, JZ, and analyzed the data, with assistance of PWL, DGB, and PH. SJR wrote the original draft of the manuscript, and all authors made substantial contributions to its form, and have critically reviewed the manuscript before submission.

