## Supplemental Figures for "The planktonic microbiome of the Great Barrier Reef"

### Supplementary Figures

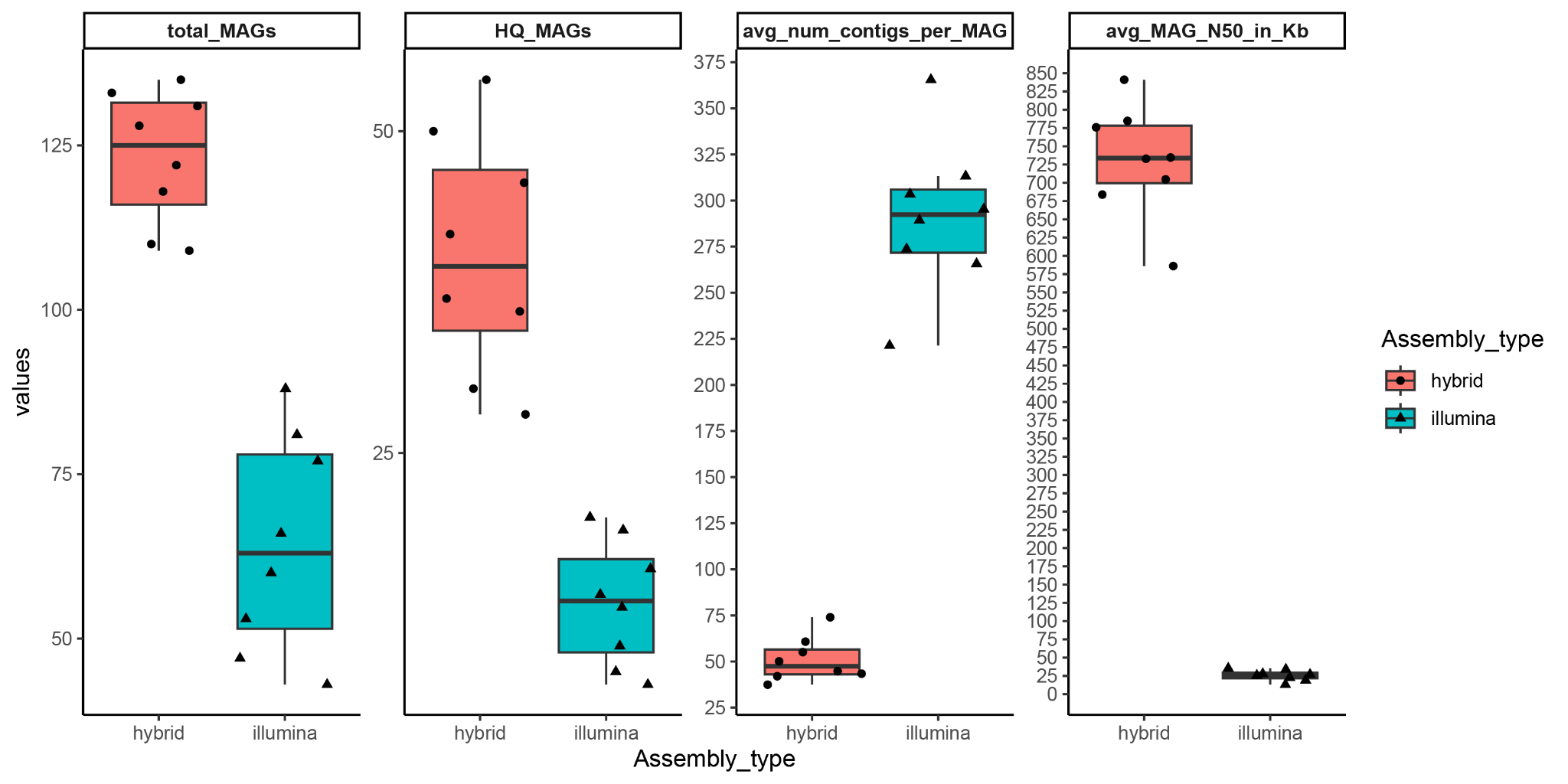

**Figure S1: MAG quality stats for Illumina-only and hybrid MAGs.** Box and whisker plot showing the number of MAGs recovered with quality >50 between Illumina-only and Nanopore Hybrid MAGs, number of high quality MAGs recovered, average number of contigs per MAG, and the average MAG N50.

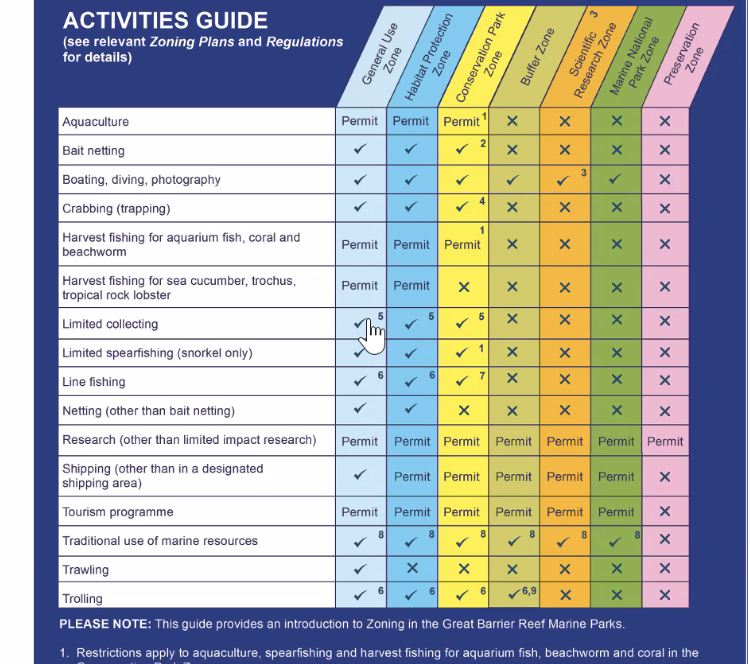

**Figure S2:** **Categories of fishing protections on the Great Barrier Reef.** In 2004, the Great Barrier Reef Marine Park Authority (GBRMPA), the government body tasked with managing the GBR, designated ~33% of the reef as open or closed to fishing, with 8 different types of zone within this scheme. In this manuscript, No-Take reefs refer exclusively to Green Marine National Park Zones that do not permit fishing in any form. In contrast, “Open” reefs here refer to either yellow Conservation Park Zones that allow Trawling for fish, or dark blue Habitat Protection Zones that allow both Trawling and Netting. We list the designation of all 48 GBR-MGD reefs in Table S8.

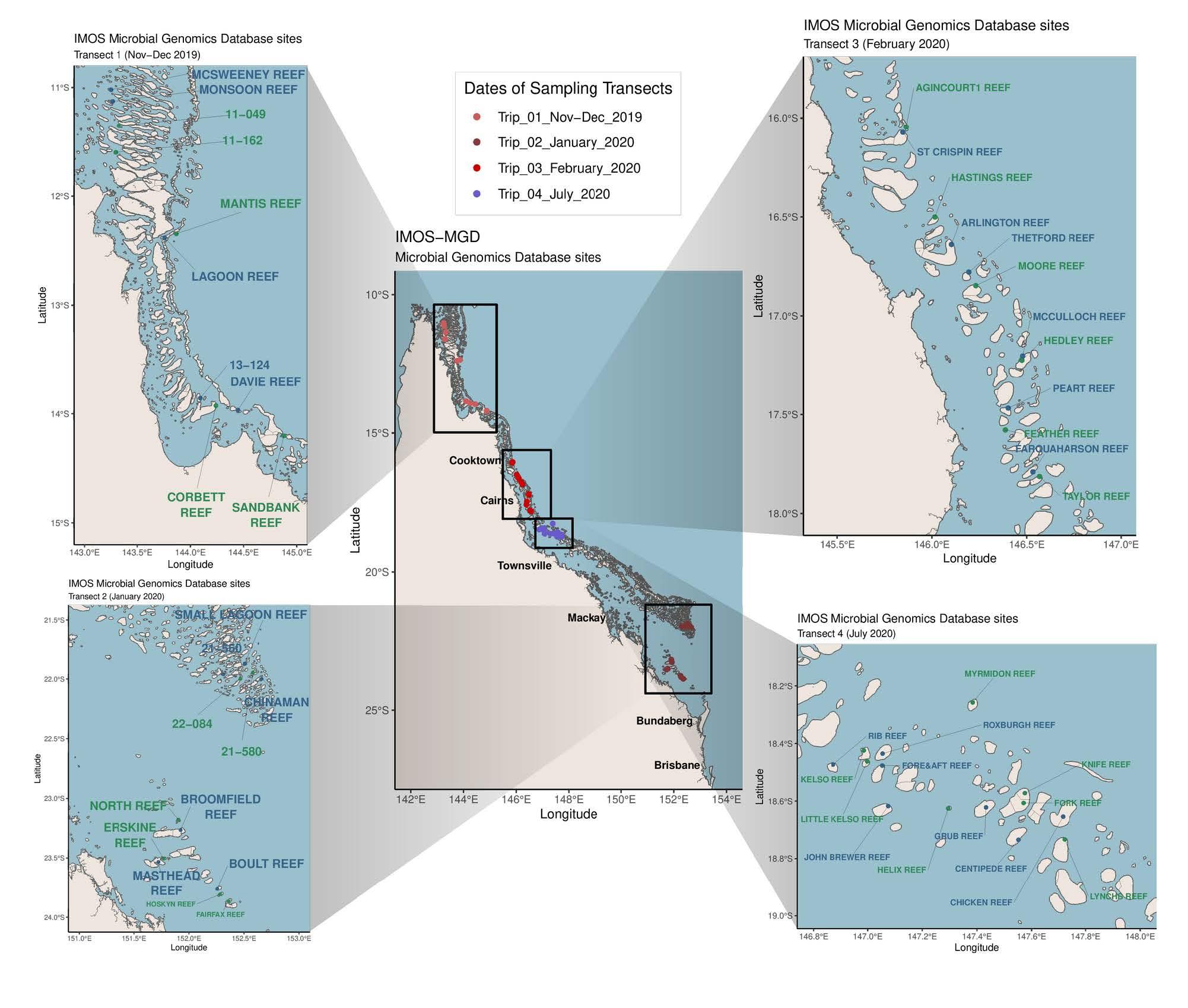

**Figure S3:** **Site map showing Fished and No-Take Marine Reserve (NTMR) reefs designated under the 2004 GBRMPA management plan.** Reefs listed in Green indicate NTMR reefs, blue indicates Fished reefs.

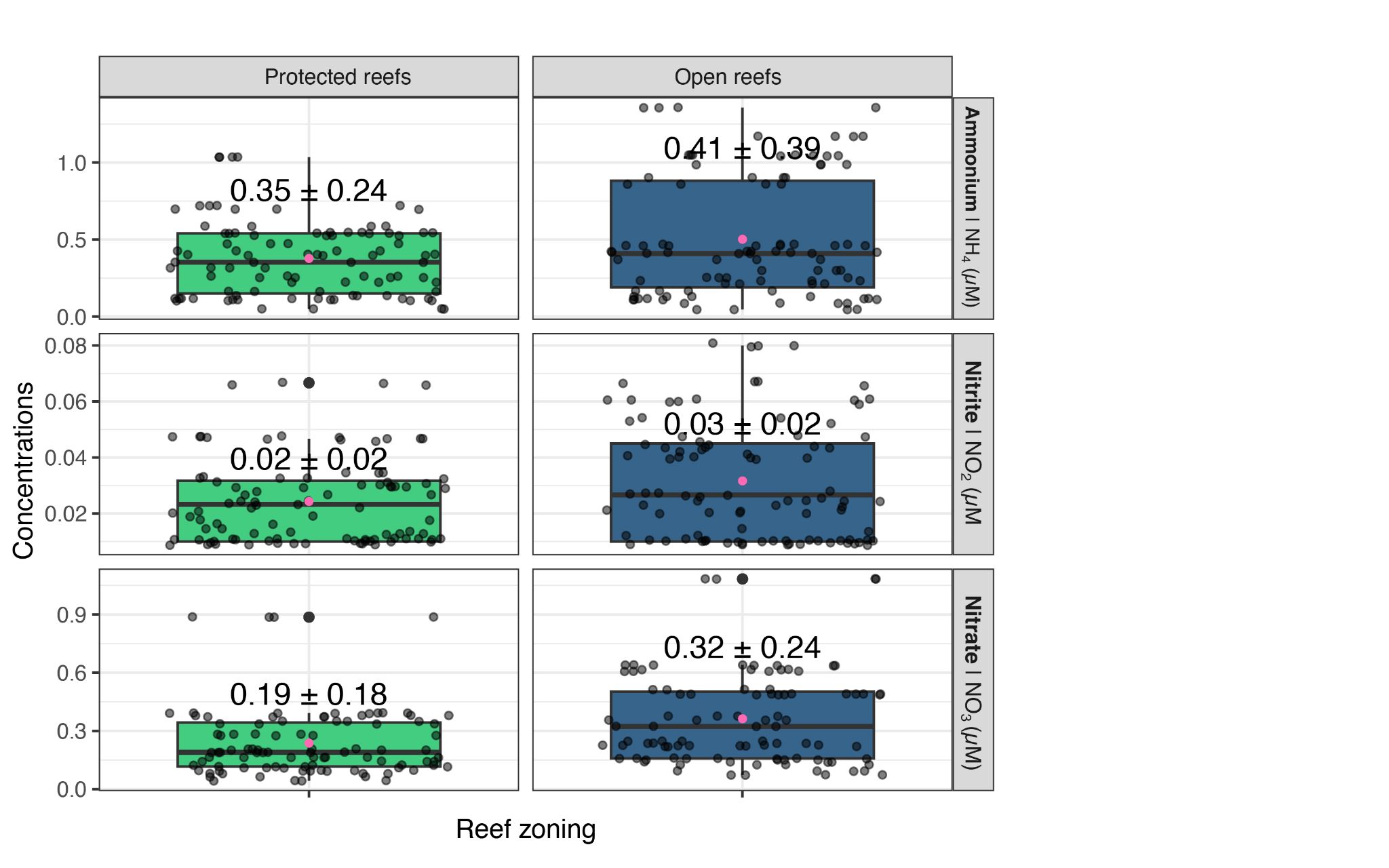

**Figure S4: Plot of dissolved nitrogen values for Fished vs No-Take Marine Reserve (NTMR) reefs.**

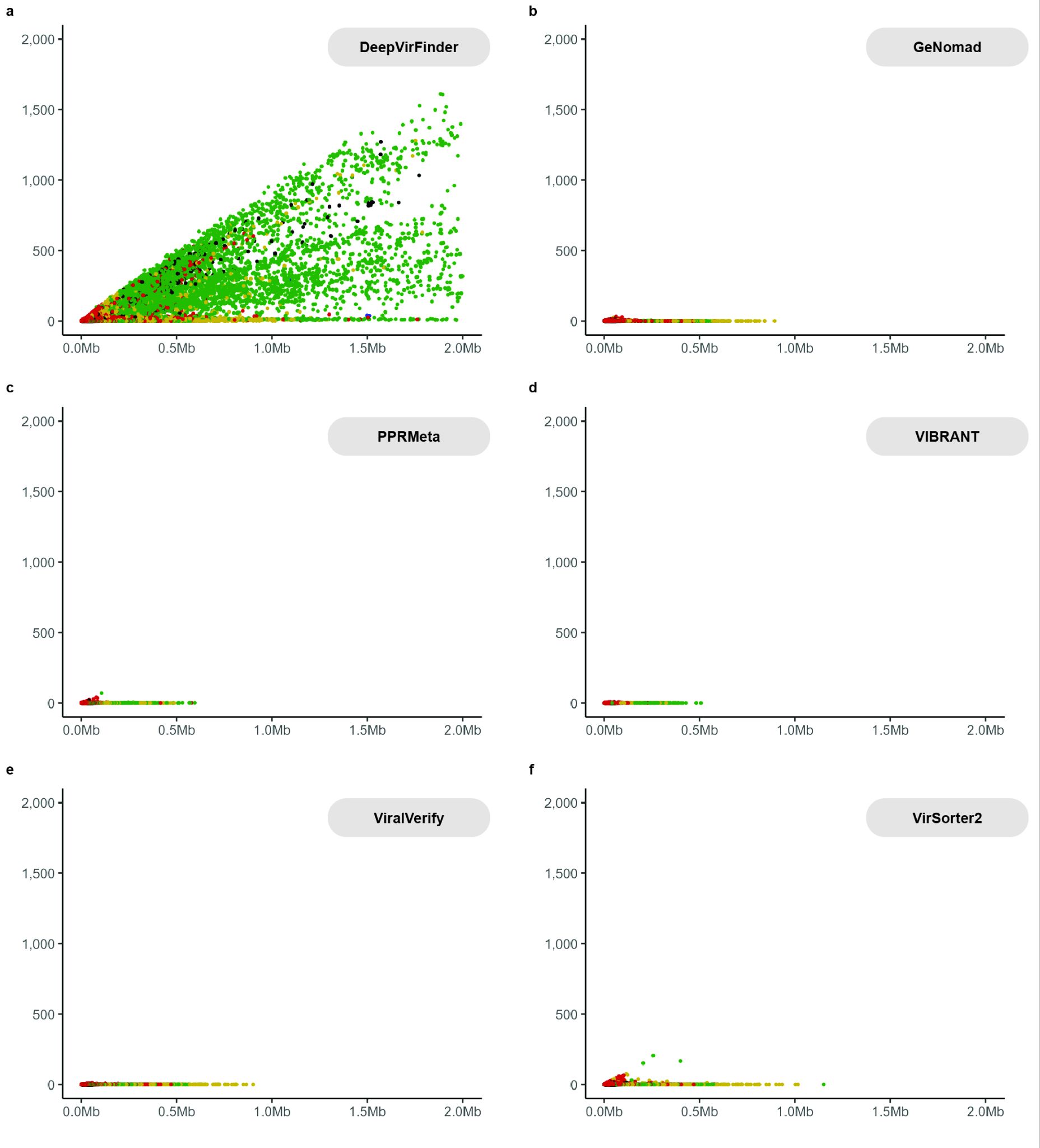

**Figure S5: Ratio of CheckV host to viral markers vs contig length in hybrid assemblies.** Dotplots show the ratio of CheckV host to viral marker genes in contigs predicted to be viral by a) DeepVirFinder, b) GeNomad, c) PPRMeta, d) VIBRANT, e) ViralVerify, and (f) VirSorter2 in hybrid Nanopore-Illumina assemblies from the GBR-MGD. Colors indicate the assigned CheckV completeness label of Complete (blue), High-Quality (green), Medium-quality (yellow), Low-Quality (red), and Not Determined (black). See Note S1 for additional discussion.

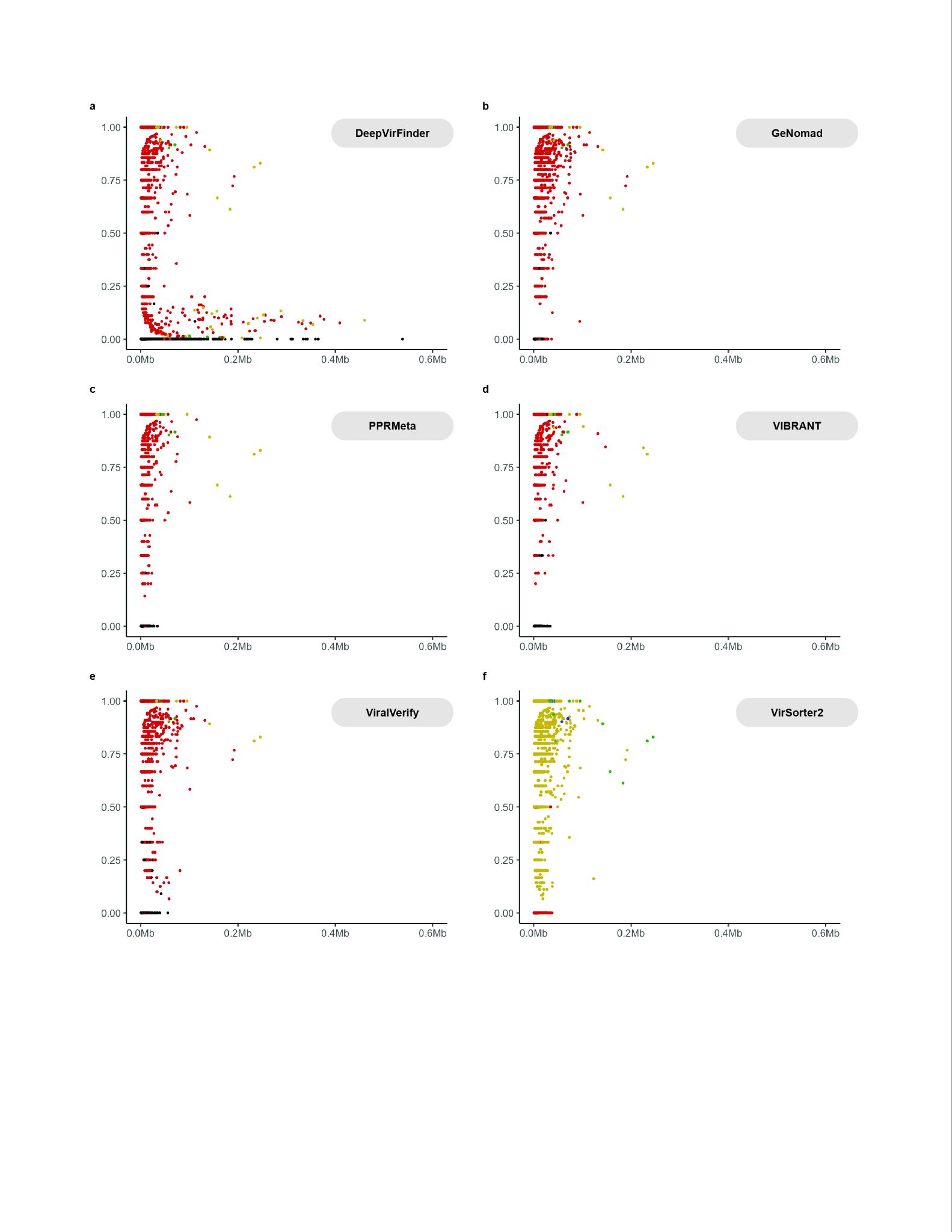

**Figure S6: Ratio of CheckV host to viral markers vs contig length in Illumina-only assemblies.** Dotplots show the ratio of CheckV host to viral marker genes in contigs predicted to be viral by a) DeepVirFinder, b) GeNomad, c) PPRMeta, d) VIBRANT, e) ViralVerify, and (f) VirSorter2 in Illumina-only assemblies from the GBR-MGD. Colors indicate the assigned CheckV completeness label of Complete (blue), High-Quality (green), Medium-quality (yellow), Low-Quality (red), and Not Determined (black). See Note S1 for additional discussion.

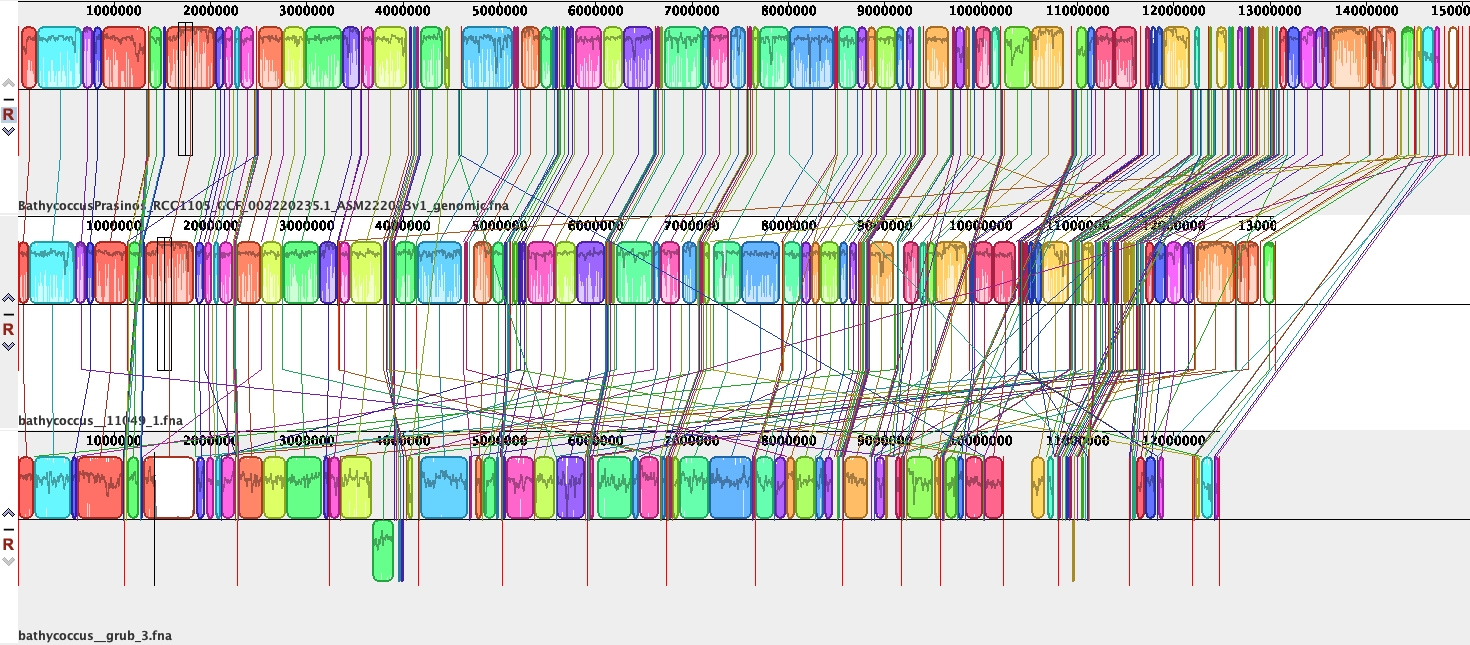

**Figure S7: Whole genome alignment of GBR-MGD Bathycoccus vs Bathycoccus prasinos RCC1105.** Alignment of the isolate reference genome from *Bathycoccus prasinos* RCC1105 (top row), *Bathycoccus prasinos* GBR_11049 from the GBR-MGD (middle), and *Bathycoccus prasinos* GBR_Grub from the GBR-MGD (bottom) using the Mauve aligner (progressiveMauve). Each chromosome is represented by one contig and Vertical red lines represent contig boundaries. The small outlier chromosome (SOC) is not shown as it is highly divergent from the reference and could not be recovered.

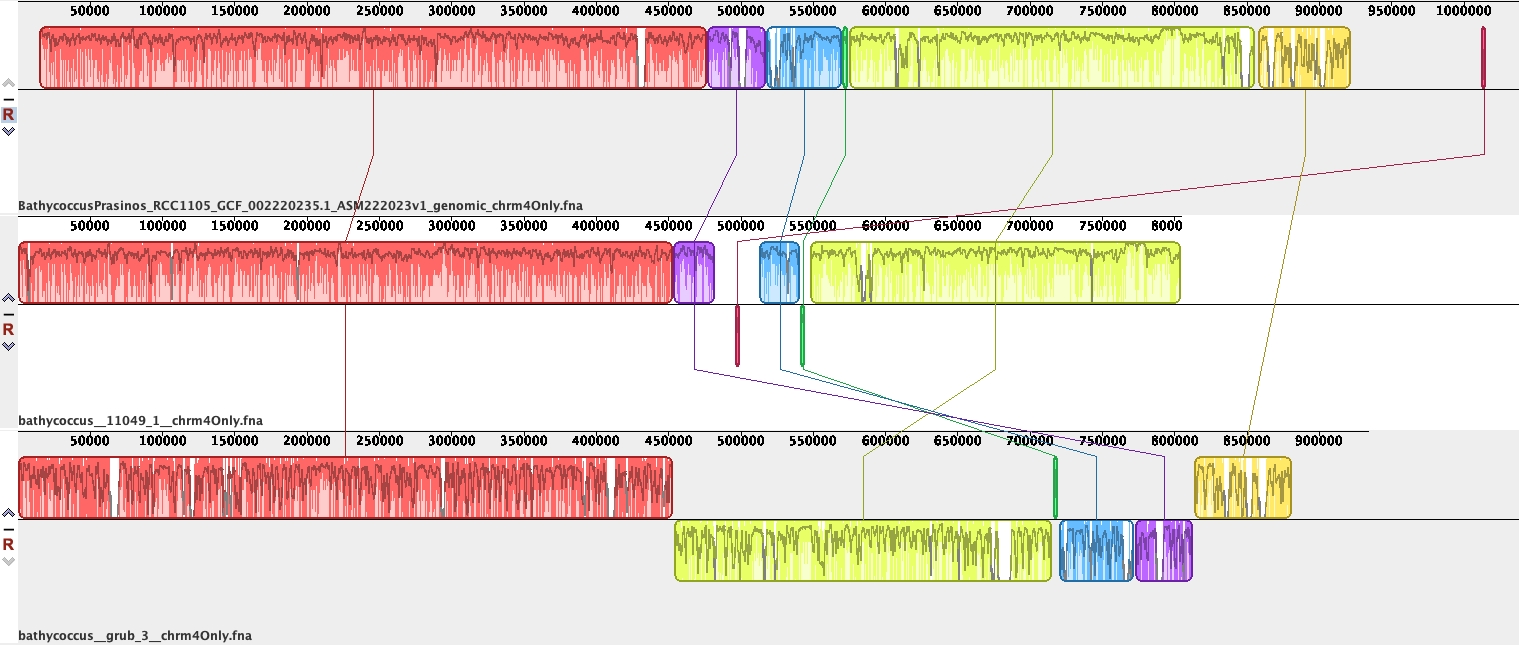
**Figure S8: Alignment chromosome 4.** Alignment of *Bathycoccus prasinos* RCC1105 (top row), *Bathycoccus prasinos* GBR_11049 from the GBR-MGD (middle), and *Bathycoccus prasinos* GBR_Grub from the GBR-MGD (bottom) using the Mauve aligner (progressiveMauve). Vertical red lines represent contig boundaries.

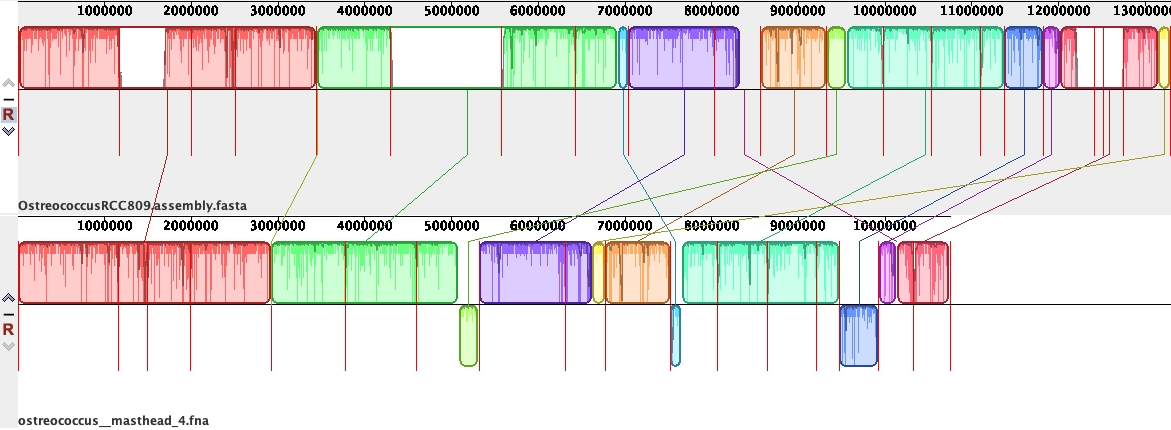

**Figure S9: Whole genome alignment of GBR-MGD *Ostreococcus* clade B vs *Ostreococcus* clade B RCC809.** Alignment of the isolate reference genome from *Ostreococcus* clade B RCC809 (top row) and *Ostreococcus* clade B from Masthead reef from the GBR-MGD (bottom) using the Mauve aligner (progressiveMauve). Each chromosome is represented by one contig and Vertical red lines represent contig boundaries.
